# Differential N-terminal processing of beta and gamma actin in vivo

**DOI:** 10.1101/2021.12.07.471626

**Authors:** Li Chen, Hsin-Yao Tang, Anna Kashina

## Abstract

Actin is one of the most essential and abundant intracellular proteins, playing an essential physiological role as the major constituent of the actin cytoskeleton. Two cytoplasmic actins, beta- and gamma-actin, are encoded by different genes, but their amino acid sequences differ only by four conservative substitutions at the N-terminus, making it very difficult to dissect their individual regulation in vivo. The majority of actins are N-terminally acetylated, following the removal of N-terminal Met. Here, we analyzed beta and gamma cytoplasmic actin N-termini in vivo and found that beta actin, unlike gamma actin, specifically undergoes sequential removal of N-terminal amino acid Asp residues. This processing affects ∼1-3% of beta actin in different cell types. We identified candidate enzymes capable of mediating this type of processing, and used CRISPR/Cas-9 to delete them, individually or together, in mammalian cell lines. This deletion abolishes most of the beta actin N-terminal processing and results in changes in F-actin levels, cell spreading, filopodia formation, and cell migration, suggesting that the beta actin processing mediated by these enzymes is physiologically important to beta actin function. We propose that selective N-terminal processing of beta actin by sequential removal of Asp contributes to differentiating the functions of non-muscle actin isoforms in vivo.

## Introduction

Actin is one of the most essential and abundant intracellular proteins, playing an essential physiological role as the major constituent of the actin cytoskeleton. In mammals, actin family is represented by six highly homologous isoforms, four that are prevalent in different types of muscle, and two non-muscle actins, beta- and gamma cytoplasmic actin [1, 2].

Beta- and gamma-actin are nearly identical at the amino acid level but are encoded by different genes that are ubiquitously expressed in every mammalian cell [3]. Their amino acid sequences differ only by four conservative substitutions at the N-terminus. At the gene level, their functions in cell migration and organism’s survival dramatically differ: beta actin’s knockout in mice results in early embryonic lethality and severe impairments of fibroblasts’ ability to migrate [4], while gamma actin’s knockout leads to much milder phenotypes [1]. Our recent work showed that this difference is defined by their nucleotide, rather than amino acid sequence [5]. Moreover, we find that the differential effects of beta and gamma actin on cell migration are defined by their nucleotide coding sequence-dependent differences in actin translation rates [6].

Actins in vivo undergo a specialized N-terminal processing, in which the N-terminal Met is acetylated and removed, and the second residue – Asp in beta actin and Glu in gamma actin – is posttranslationally acetylated [7]. Muscle actins undergo similar processing, even though differences in their N-termini account for slightly different steps preceding acetylation in muscle and non-muslce cells [8]. Overall, >90% of intracellular actin is acetylated in steady state, and this acetylation facilitates actin’s cellular functions [9].

Actin undergoes multiple posttranslational modifications (PTM) [10, 11], however most of these modifications are believed to uniformly affect all actin isoforms, and any examples to the contrary are very difficult to parse out due to the near-identity of actin amino acid sequences and their coexistence in the same cells. One known exception constitutes N-terminal arginylation, which targets the only sequence sufficiently different between non-muscle actins to detect by the currently available methods. Earlier studies from our group found that beta, but not gamma actin undergoes N-terminal arginylation on Asp 3, presumably exposed after a previously unknown N-terminal processing step involving removal of the second Asp [12]. Such N-terminal beta actin arginylation affects ∼0.1% of the total intracellular beta actin pool [13], making it extremely difficult to detect. Our work suggest that selective arginylation of beta, but not gamma, actin is defined by their nucleotide coding sequence [14], likely linked to their different translation rates [6], and that this arginylation is essential for cell migration [12]. However, processing steps leading to beta actin’s arginylation on Asp 3 has never been described.

Here, we analyzed beta and gamma cytoplasmic actin N-termini in vivo and found that beta actin undergoes sequential removal of N-terminal amino acid Asp residues, resulting in intracellular actin species lacking one, two, or three Asp from the N-terminus. In contrast, gamma actin does not appear to undergo such processing, and only the N-termini containing Met, or acetylated Asp after Met removal, can be seen. Overall, this processing affects ∼1-3% of the total beta actin pool in different cell types. We identified two candidate enzymes, N-terminal Asp and Glu aminopeptidases (DNPEP and ENPEP), and deleted them in human HAP1 cells to assess their role in beta actin processing and the basic properties of the actin cytoskeleton-dependent intracellular processes. We find that while deletion of *DNPEP* does not strongly affect removal of beta actin’s N-terminal Asp, *ENPEP* deletion has a much more pronounced effect, and the double knockout leads to nearly complete abolishment of this processing. *DNPEP* and *ENPEP* knockout cells display differences in F-actin levels, cell spreading, filopodia formation, and cell migration, suggesting that the beta actin processing mediated by these enzymes is physiologically important for beta actin function. In vitro experiments with N-terminal beta and gamma actin derived peptides show that ENPEP is capable of targeting both unprocessed beta- and gamma-actin N-termini after Met removal, but neither ENPEP or DNPEP can target the peptides containing N-terminally acetylated Asp and Glu, suggesting that in vivo targeting of this processing to beta actin requires additional recognition mechanisms, and that this processing likely occurs prior to actin acetylation. We propose that selective N-terminal processing of beta actin by sequential removal of Asp contributes to differentiating the functions of non-muscle actin isoforms in vivo.

## Results

### Beta and gamma actin are differentially N-terminally processed in vivo

To analyze the processing of the actin’s N-termini in non-muscle cells, we performed mass spectrometry of total intracellular actin from mouse embryonic fibroblasts (MEF) cell, a commonly used non-muscle migratory cell type that expresses approximately equal levels of beta and gamma cytoplasmic actin.

To capture an enriched fraction of total actin without any bias in purification steps, we fractionated total cell lysates on SDS page and broadly excised a prominent protein band in the ∼43 kDa range corresponding to the actin’s molecular weight (Fig. S1). We then performed in-gel digestion of this band by trypsin and analyzed it by LC-MS/MS. To ensure identification of the processed actin N-termini, we performed searches under semi-tryptic conditions, since fully tryptic search can only identify unprocessed or minimally processed N-termini. We also included the most common N-terminal modifications, including N-terminal acetylation abd Met and Cys oxidation, which can occur both natively and during sample preparation.

This analysis revealed a repertoire of N-terminal peptides derived from both beta and gamma actin (Fig. 1). As previously shown, the majority of these peptides (>95%) represented N-terminal acetylation on Asp 2 or Glu 2 in beta and gamma actin, respectively (Fig. 1, top left). A small fraction of the peptides were unprocessed, containing N-terminal initiator Met. Curiously, the fraction of Met containing peptides was somewhat higher in gamma actin (∼3%) compared to beta actin (∼1%) (Fig. 1, top left). This suggests that the enzyme(s) mediating this removal have a slight preference for beta actin.

**Figure 1.**
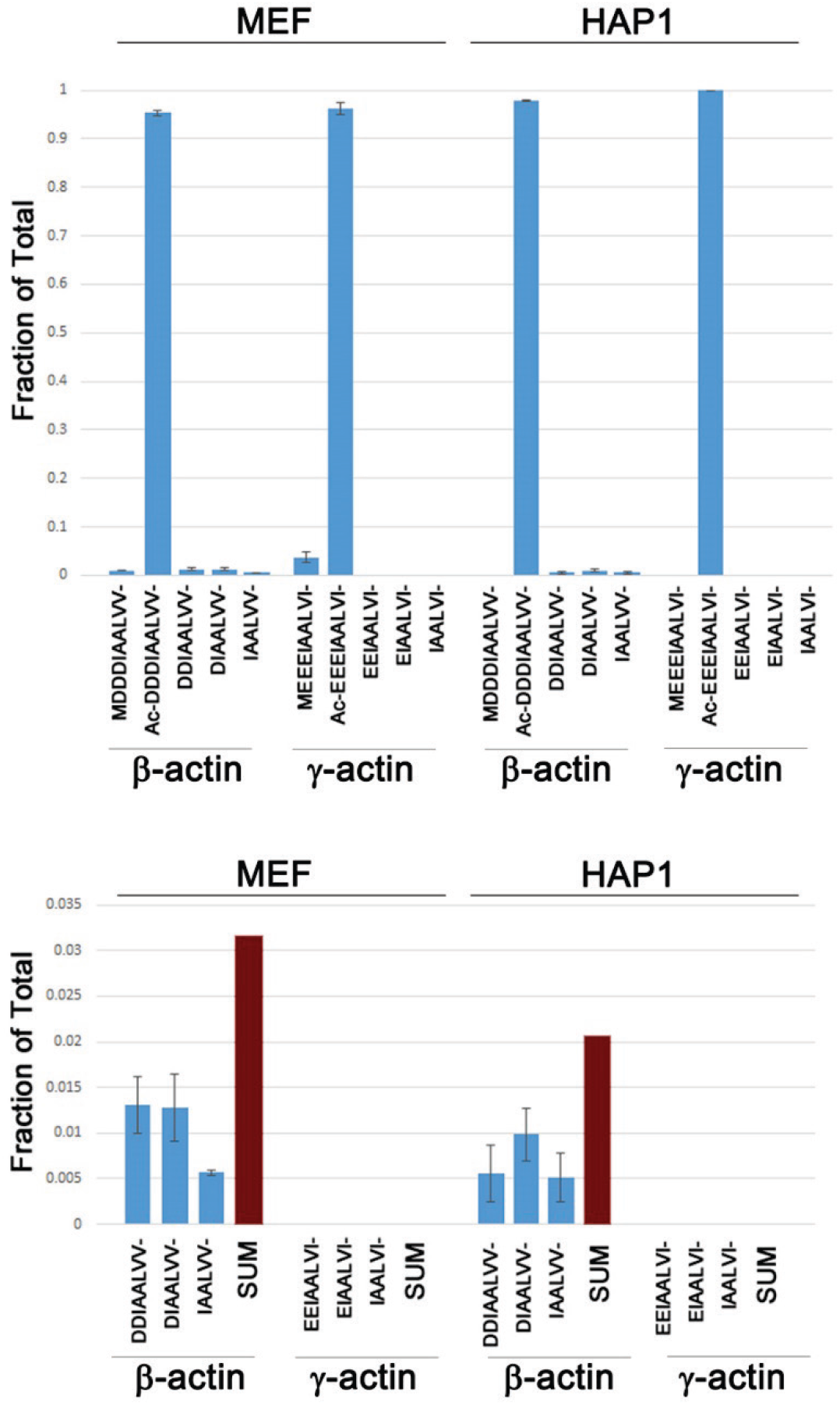
Beta and gamma actin are differentially N-terminally processed in mammalian cells. Mass spectrometry quantification of the N-terminal beta- and gamma-actin peptides in mouse embryonic fibroblasts (MEF) and human HAP1 cells, quantified as the ratio of intensities for each peptide versus the total intensities for the entire set of N-terminal peptides identified for each actin isoform (fraction of the total). N-terminal sequence of each peptide for each isoform is indicated on the x-axis. Upper panel, the entire repertoire of N-terminal peptides found in cells. Lower panel, expanded view of the peptides generated by sequential removal of N-terminal Asp residues. Error bars represent SEM, n=3.

The variety of peptides derived from the beta actin’s N-termini was greater than that of gamma actin. In addition to Met-Asp-Asp-Asp-Ile-Ala-Ala-Leu-(MDDDIAAL-) and Acetyl-Asp-Asp-Asp-Ile-Ala-Ala-Leu-((Ac-DDDIAAL-) N-termini, we also detected the N-terminal peptides sequentially lacking each of the Asp (DDIAAL-, DIAAL-, and IAAL-). Overall, these processed actin species accounted for ∼3% of the total beta actin (Fig. 1, bottom left). No such peptides were detected for gamma actin, which existed solely as a mixture of unprocessed and N-terminally acetylated species (Fig. 1, left).

To test whether similar beta actin processing also occurs in other cell types, we analyzed HAP1 cells, a near-haploid adherent fibroblast-like cell line of human leukemia origin [15] commonly used in CRISPR/Cas9 knockout studies. In these cells, no Met containing actin species were detected, suggesting that the removal of N-terminal Met occurs much more rapidly and efficiently than in MEFs. 100% of gamma actin existed as acetylated species (Fig. 1, top right). However, beta actin in these cells also contained a smaller but detectable fraction of peptides that had undergone sequential removal of Asp 2, Asp 3, and Asp 4. This processing affected ∼1% of the total beta actin (Fig. 1, bottom left).

Thus, sequential removal of N-terminal Asp residues is specific to beta actin and can occur in different cell types.

### Differential N-terminal processing of beta and gamma actin is defined by their amino acid sequence

We previously found that beta actin’s key functions in cell migration and organism’s survival are defined by its nucleotide, rather than amino acid sequence [6]. We also found that N-terminal arginylation of beta actin is defined by its nucleotide coding sequence and translation rate [15]. To test whether the sequential removal of N-terminal Asp residues in beta actin is nucleotide coding sequence dependent, we used mass spectrometry to analyze MEF in which the endogenous beta actin gene has been edited using CRISPR/Cas9 to encode gamma actin protein, without substantially affecting the nucleotide sequence (beta coded gamma actin or ACTBcG [5]). In these cells, gamma actin protein is expressed from both the endogenous beta- and gamma-actin genes. Thus, if the differential N-terminal processing of beta and gamma actin is nucleotide sequence-dependent, N-terminally processed gamma actin with sequentially removed N-terminal Glu should be detected in these cells, unlike in control. However, we found no evidence of such processing (Fig. 2). In both wild type and ACTBcG cells, the only gamma actin species we detected were either unprocessed, containing N-terminal Met, or acetylated on Glu 2, which constituted the majority of the pool (Fig. 2, left). Curiously, the fraction of Met-uncleaved gamma actin in ACTBcG cells was lower than in control (Fig. 2, right), consistent with the observation that beta actin’s N-terminal Met removal appears more efficient than gamma actin (Fig. 1, top left). Thus, this difference in Met removal between beta and gamma actin appears to be nucleotide coding sequence dependent. However, further processing of actin by sequential removal of second, third, and fourth negatively charged residues is clearly dependent on the amino acid sequence.

**Figure 2.**
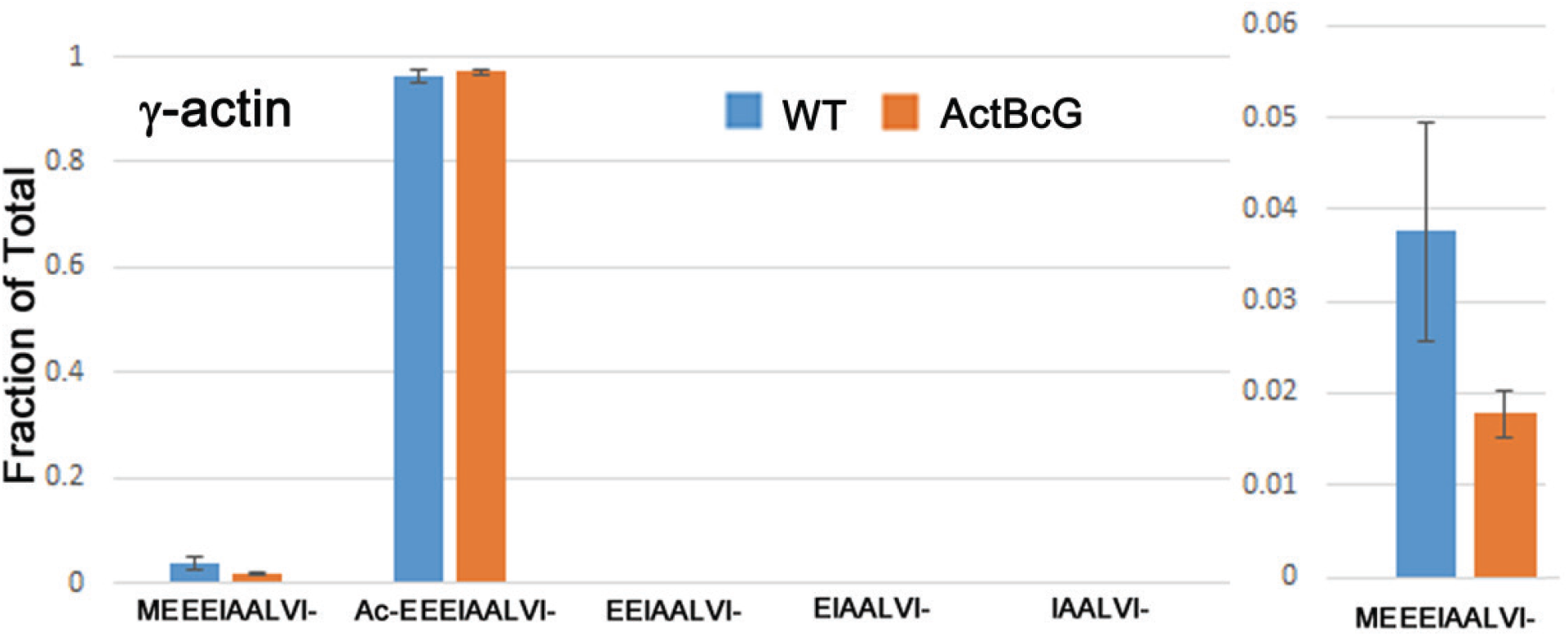
Lack of sequential removal of N-terminal amino acid residues in gamma actin is not nucleotide coding sequence-dependent. Mass spectrometry quantification of the N-terminal gamma-actin peptides in wild type (WT) and gene-edited MEFs expressing gamma actin protein sequence from both beta and gamma actin gene (ActBcG). Ratios of intensities for each peptide versus the total intensities for the entire set of N-terminal peptides are shown on the y-axis (fraction of the total). N-terminal sequence of each peptide for each isoform is indicated on the x-axis. Error bars represent SEM, n=3.

### Deletion of N-terminal Glu aminopeptidase greatly inhibits sequential removal of N-terminal residues from beta actin in vivo

We searched the mouse genome for the candidate enzymes that could perform this processing of the actin’s N-termini. The only two aminopeptidases previously shown to possess the activity to remove N-terminal Asp or Glu were Aspartyl and Glutamyl Aminopeptidases (DNPEP and ENPEP). Both enzymes are capable of removal of negatively charged residues from the peptide/protein N-termini ([16, 17]) and do not appear to strongly discriminate between Asp and Glu. Thus, we used CRISPR/Cas9 to knock out both *DNPEP* and *ENPEP*, individually or together, in HAP1 cells. We then analyzed the effect of these knockouts on the actin N-terminal processing, using three independently derived cell lines from each genotype (Fig. 3).

**Figure 3.**
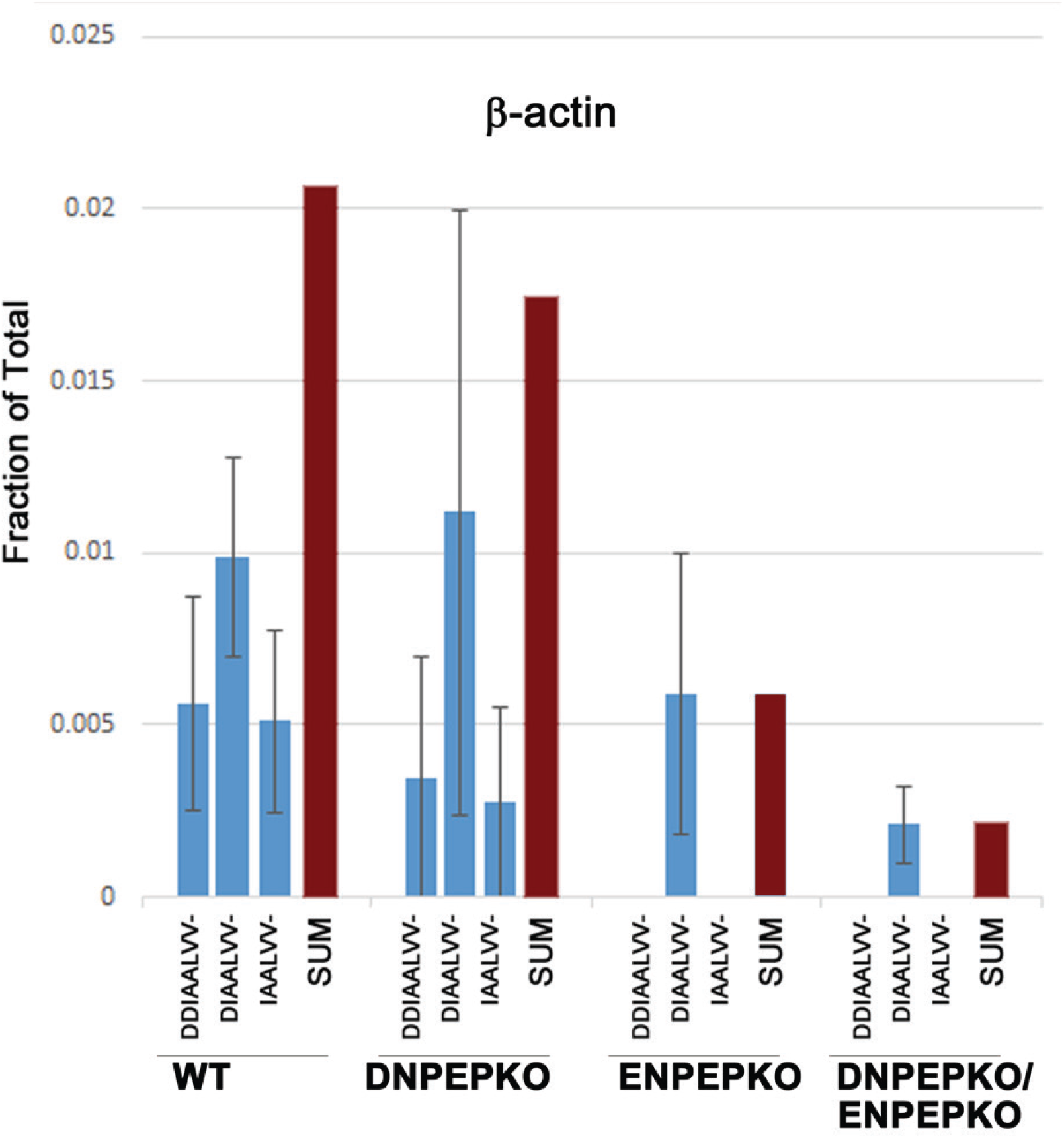
Deletion of Aspartyl and Glutamyl Aminopetidases DNPEP and ENPEP inhibits sequential removal of acidic amino acid residues from beta actin N-terminus. Quantification of N-terminal processing of beta actin from WT, *DNPEP*-KO, *ENPEP*-KO and double knockout HAP1 cells, presented as fraction of the total similarly to that described in Fig. 1 and 2. Error bars represent SEM, n=3 independently derived cell lines for each genotype.

Knockout of *DNPEP* did not strongly affect the N-terminal processing of beta actin. In two out of three knockout lines, we did not detect any peptides with DDIAAL- or IAAL-N-terminal sequence, and one out of three did not contain DIAAL-peptides, however between the three replicates, at least one type of processing was observed in each, suggesting that this knockout might reduce, but not abolish the processing. *ENPEP* knockout had a much stronger effect, with no DDIAAL- or IAAL-peptides detected in any replicate cell lines and the amount of DIAAL-peptide greatly reduced. Double knockout *DNPEP*/*ENPEP* had only trace amount of DIAAL-peptide, and no other peptides from these processing steps detected. Thus, it appears that ENPEP accounts for the majority of the processing, and DNPEP potentially contributes to it, even though it is possible that the DIAAL-peptide can also be generated by some additional enzymatic machinery.

### N-terminal Glu aminopeptidase targets non-acetylated actin N-termini and does not differentiate between actin isoforms in vitro

Both DNPEP and ENPEP have been characterized as the enzymes that can target N-terminal acidic residues, without an apparent bias toward N-terminal Asp or Glu. With this knowledge, it is unclear how these two enzymes can distinguish between beta and gamma actin, both of which contain a stretch of three acidic residues at the N-terminus. We used in vitro assays with synthetic peptides corresponding to the beta and gamma actin’s N-terminal sequence to assess whether these enzymes exhibit any specificity toward one of the actin isoforms. In these in vitro assays DNPEP did not yield any detectable N-terminally processed peptides, suggesting that this enzyme’s activity is low, or actin’s N-terminal peptides are not its preferred substrates (Fig. 4 left, Fig. S2A). Only ENPEP was able to mediate sequential removal of acidic residues from the peptides’ N-termini, with all the processed peptide variants detected after the reaction (Fig. 4 right, Fig. S2B). However, this enzyme did not distinguish between beta and gamma actin’s N-terminal peptides in vitro and processed both with apparently equal efficiency. Thus, if ENPEP accounts for the majority of beta actin’s N-terminal processing, as suggested by our in vivo data, this enzyme’s specificity toward beta rather than gamma actin in vivo is likely mediated by additional factors or physiological processes that counteract the action of these enzymes in the context of actin isoforms.

**Figure 4.**
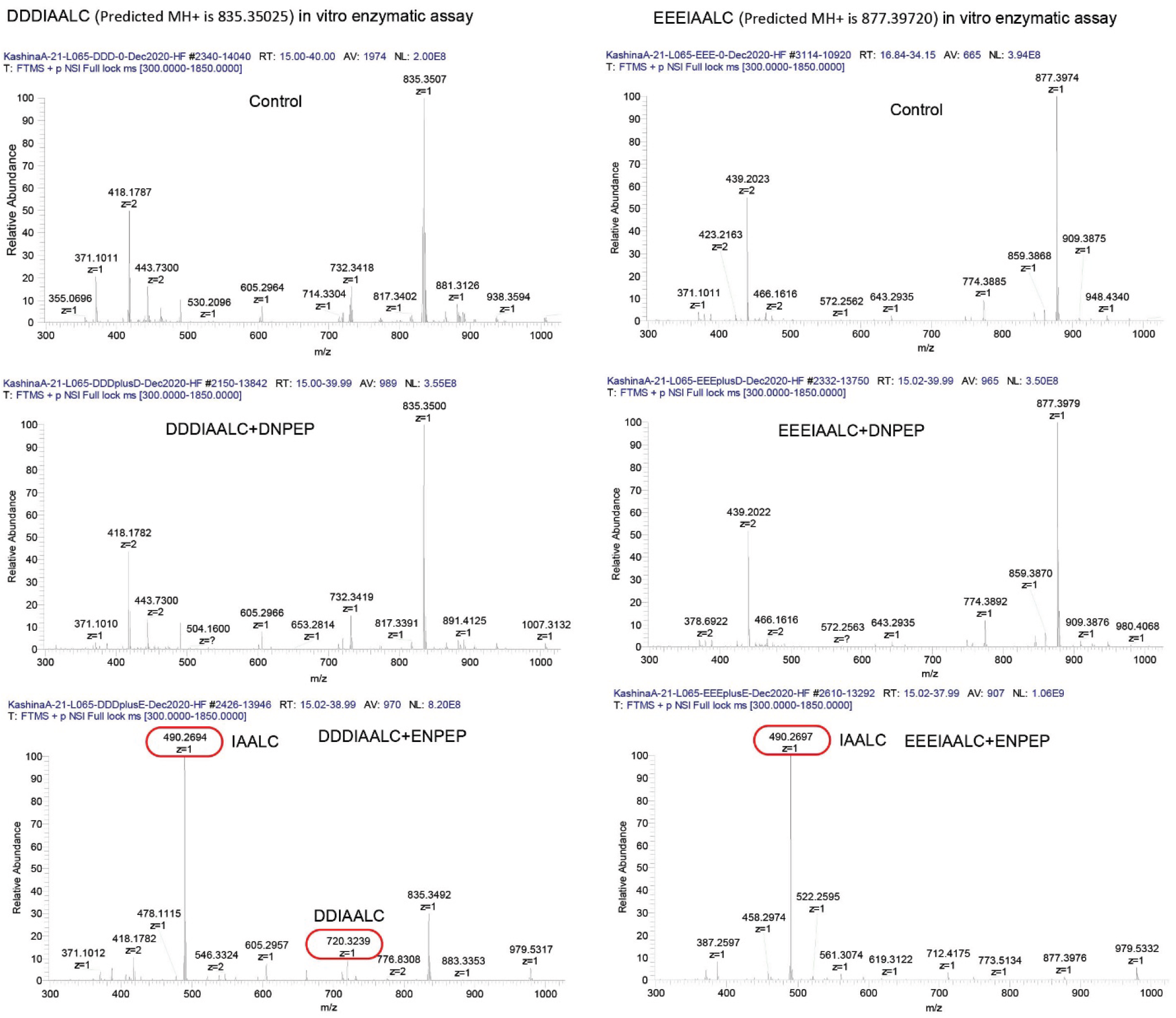
Glutamyl aminopeptidase ENPEP can process N-termini of beta and gamma actin in vitro. Mass spectrometry analysis of beta and gamma actin N-terminal peptides before (control) and after incubation with purified ENPEP and DNPEP. Red frames indicate the enzymatically processed peptides. ENPEP, but not DNPEP, is able to generate peptides lacking one or more N-terminal acidic residues in both beta (left) and gamma (right) actin-like peptides

Most of the actin in vivo is N-terminally acetylated ([9] and Fig. 1). To test whether DNPEP and ENPEP can act on the N-terminally acetylated actin peptides, we performed in vitro assays with synthetic peptides containing an acetyl group to mimic the native sequence of beta and gamma actin. In this assay, neither of the enzymes were able to process the acetylated N-terminal Asp of Glu in the beta- or gamma-actin sequence (Fig. 5, Fig. S3). This result indirectly suggests that the sequential processing of the actin N-terminal residues by DNPEP and/or ENPEP occurs on non-acetylated actin, likely prior to its targeting by actin N-terminal acetyltransferase NAA80.

**Figure 5.**
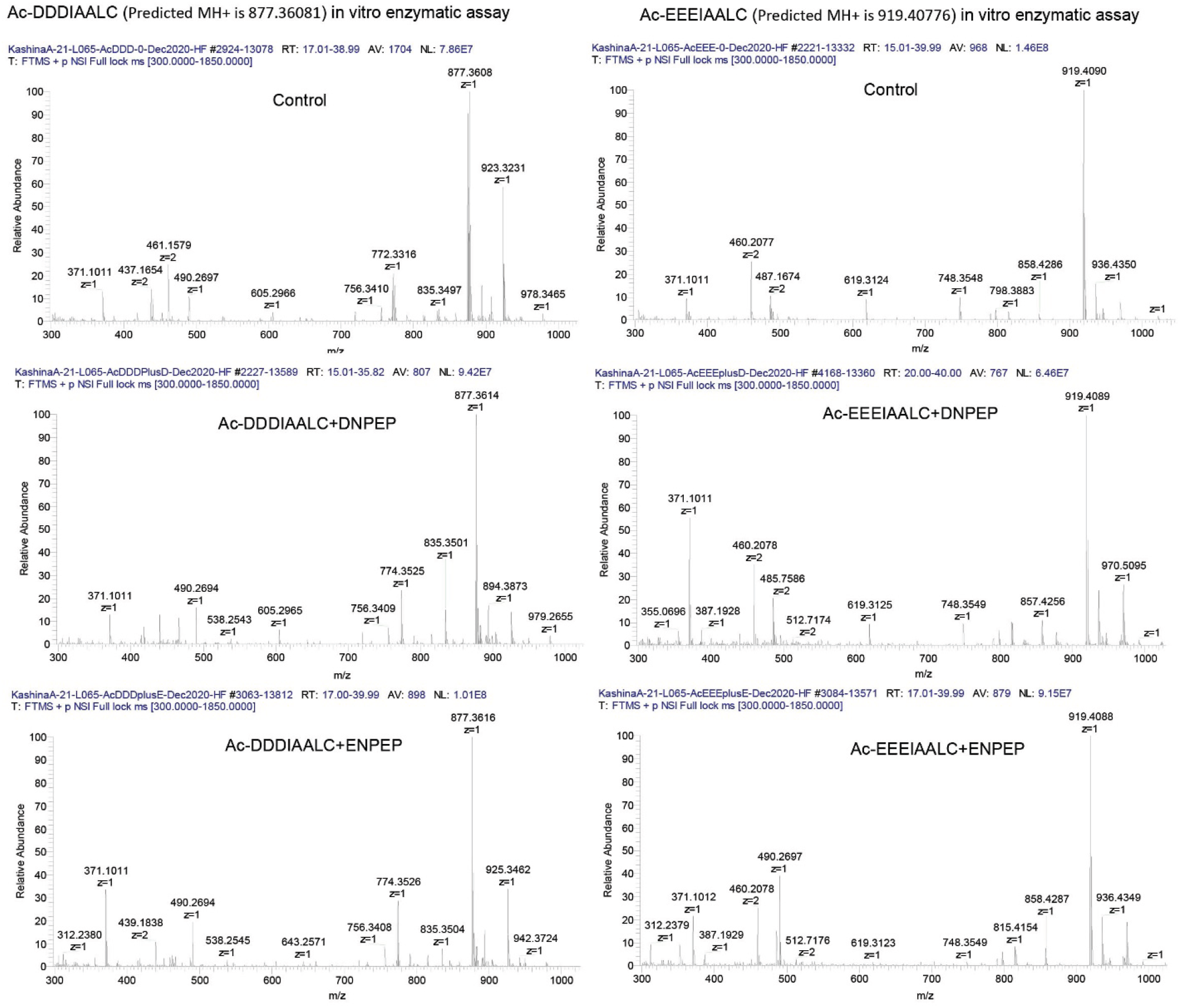
Aspartyl and Glutamyl aminopeptidases DNPEP and ENPEP are unable to process acetylated N-termini of beta and gamma actin in vitro. Mass spectrometry analysis of acetylated beta and gamma actin N-terminal peptides before (control) and after incubation with purified ENPEP and DNPEP. No processing is observed with either peptide, suggesting that N-terminal acetylation prevents removal of acidic N-terminal residues.

### Deletion of N-terminal Asp and Glu aminopeptidase genes affects actin cytoskeleton, filopodia formation, and cell spreading

To test whether DNPEP and ENPEP contribute to actin cytoskeleton-related processes in vivo, we analyzed the morphology and basic actin cytoskeleton-dependent processes in *DNPEP* and *ENPEP* knockout HAP1 cells. First, we measured the overall F-actin levels by comparing the average and total staining with fluorescent phalloidin per cell. The unit area intensity was slightly higher in *DNPEP* and *DNPEP/ENPEP* double knockout, suggesting that DNPEP is required for maintaining spatial arrangement of the actin cytoskeleton at the total cell level (Fig. 6). This parameter appeared to be unchanged in the *ENPEP* knockout compared to control. In contrast, the total cell intensity of fluorescent phalloidin was significantly lower in *ENPEP* knockout compared to WT, suggesting that this knockout leads to an overall reduction in the amount of polymerized actin per cell. No differences in overall F-actin levels per cell were observed in *DNPEP* knockout cells, and *DNPEP/ENPEP* double knockout showed the same trend as *ENPEP* knockout but with higher p value, possibly due to greater variabilities between replicates in this cell line (Fig. 6). Thus, *ENPEP* knockout results in an overall reduction of polymeric actin and *DNPEP* knockout results in potential spacial rearrangement of the actin cytoskeleton.

**Figure 6.**
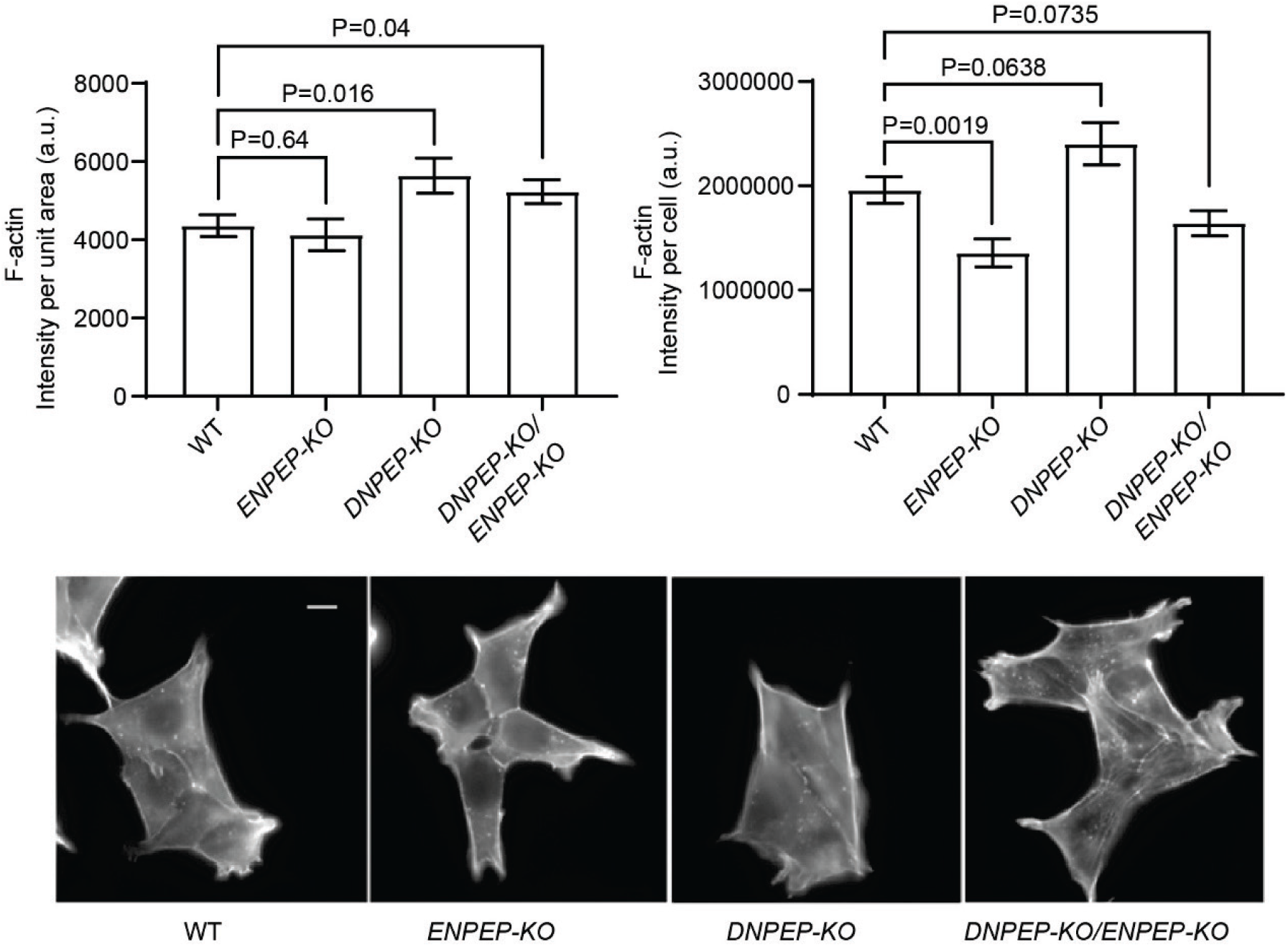
DNPEP and ENPEP knockout results in changes in actin polymer levels. Top, quantification of F-actin per unit area (left) and F-actin per cell (right) detected by Phalloidin-AlexaFluor488 staining of different genotypes; bottom, representative images of different genotypes stained with Phalloidin-AlexaFluor488 (scale bar, 10 µm). Error bars represent SEM, n= 31 for WT, 30 for ENPEPKO, 28 for DNPEP KO, and 31 for the double knockout for the intensity per cell measurement, and 31 for WT, 30 for ENPEPKO, 29 for DNPEP KO, and 32 for the double knockout for the intensity per unit area measurement.

The number and length of filopodia in both *DNPEP* and *ENPEP* knockout cells was higher than in wild type, and this effect was more pronounced in the double knockout (Fig. 7), suggesting that both enzymes contribute to this process in a cumulative manner.

**Figure 7.**
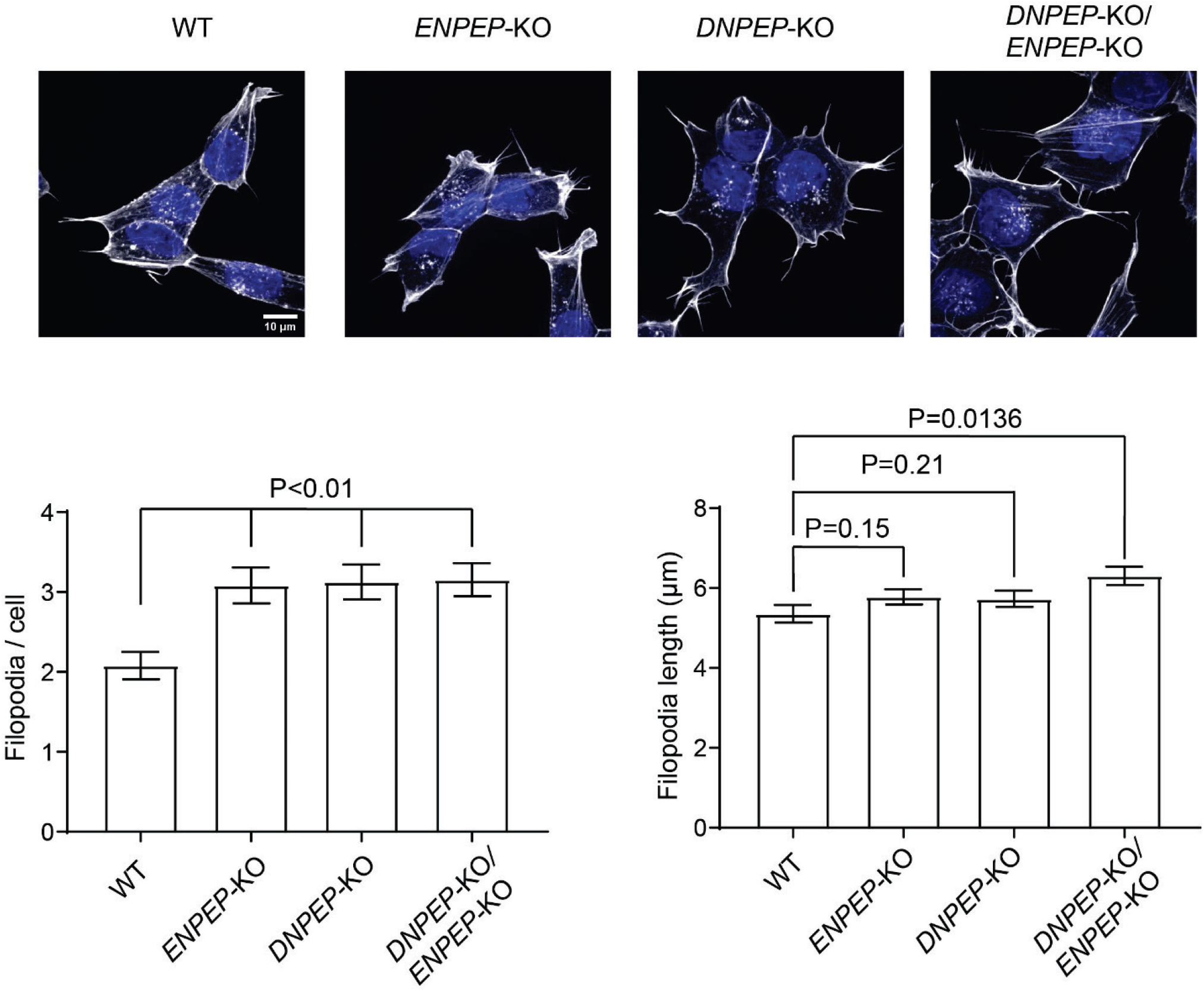
DNPEP and ENPEP knockout results in changes in the number and length of filopodia. Top, Representative images of cells from each genotype stained with Phalloidin-AlexaFluor488 (scale bar, 10 µm); bottom, filopodia number (left) and length (right) quantification. Error bars represent SEM, n=181 for WT, 259 for ENPEPKO, 222 for DNPEP KO, and 205 for the double knockout for the filopodia length measurement, and 84 for WT, 84 for ENPEPKO, 71 for DNPEP KO, and 65 for the double knockout for the filopodia numbers measurement.

*ENPEP* knockout, as well as *DNPEP*/*ENPEP* double knockout, resulted in a significant and pronounced decrease in cell spreading (measured as the area occupied by cells spread on the substrate, Fig. 8, left). This effect was not due to a reduction in cell volume, since the size of the spherical trypsinized cells was similar in all cell types (Fig. 8, right). Notably, no reduction in cell spreading was observed in *DNPEP* knockout, suggesting that ENPEP is the sole enzyme responsible for this effect.

**Figure 8.**
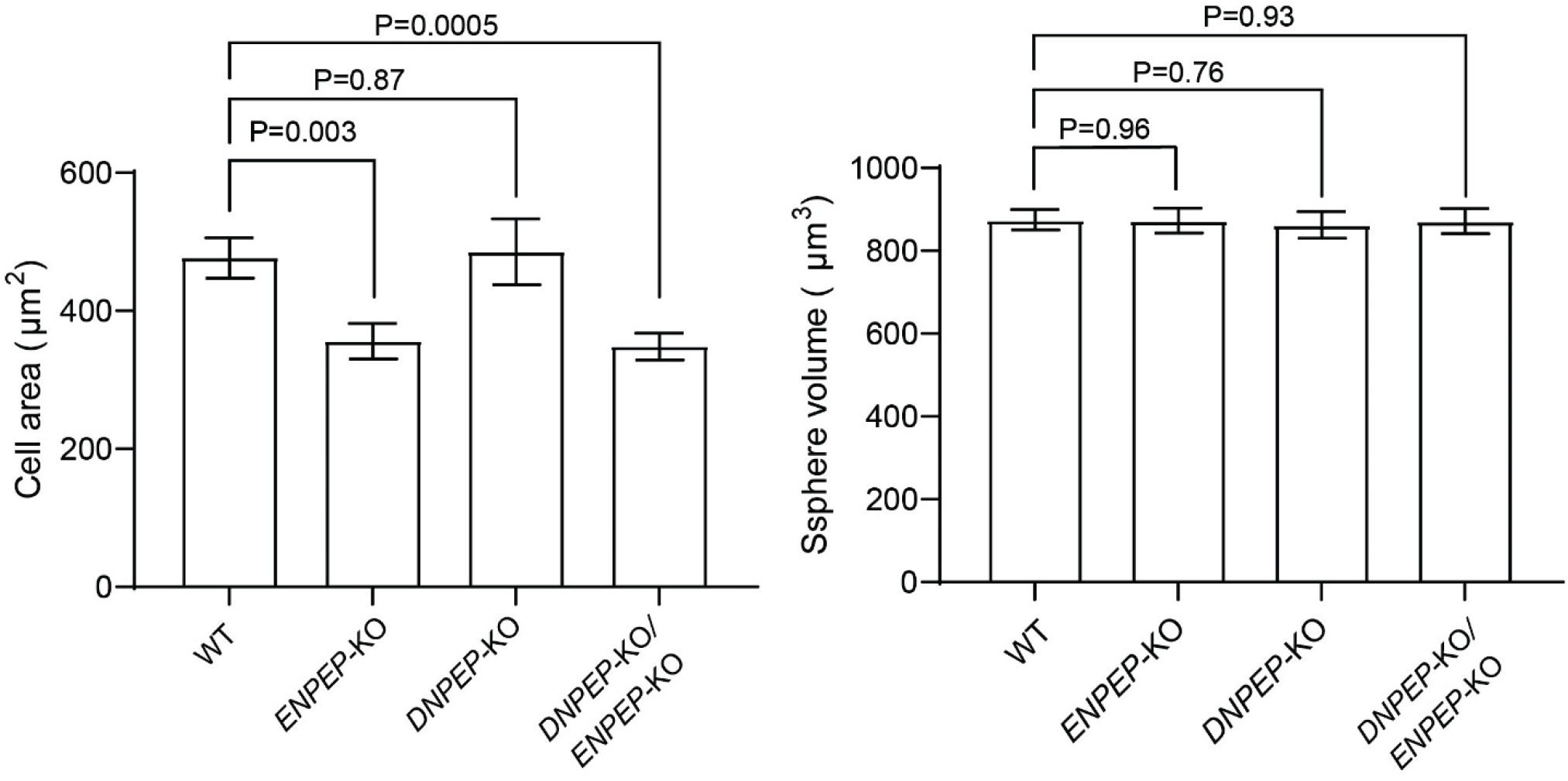
ENPEP knockout results in reduced cell spreading. Left, quantification of the area occupied by spread cells on the substrate; right, quantification of cell volume, derived from the measurements of trypsinized spherical cells in suspension. Error bars represent SEM, n= 31 for WT, 30 for ENPEPKO, 29 for DNPEP KO, and 32 for the double knockout for the cell area measurement, and 149 for WT, 156 for ENPEPKO, 166 for DNPEP KO, and 139 for the double knockout for the cell volume measurement.

Thus, both DNPEP and ENPEP contribute to actin cytoskeleton maintenance, filopodia formation and cell spreading, and differentially affect these processes, suggesting that they may play different, partially overlapping roles in the actin cytoskeleton maintenance and function.

### Deletion of N-terminal Asp and Glu aminopeptidase genes affect cell migration

One of the key functions of non-muscle actin cytoskeleton is its role in cell migration, and it has been previously found that in non-muscle cells beta actin plays a critical role in this process [18]. We therefore compared the rates of migration of HAP1 cells of all four genotypes into an infinite scratch wound in cell culture (Fig. 9). *ENPEP* knockout did not substantially affect cell migration rates, however, but *DNPEP* knockout resulted in substantially slower migration, and this effect was even more pronounced in the double knockout cells. Thus, *DNPEP* critically affects cell migration, even though *ENPEP* potentially also contributes to this process.

**Figure 9.**
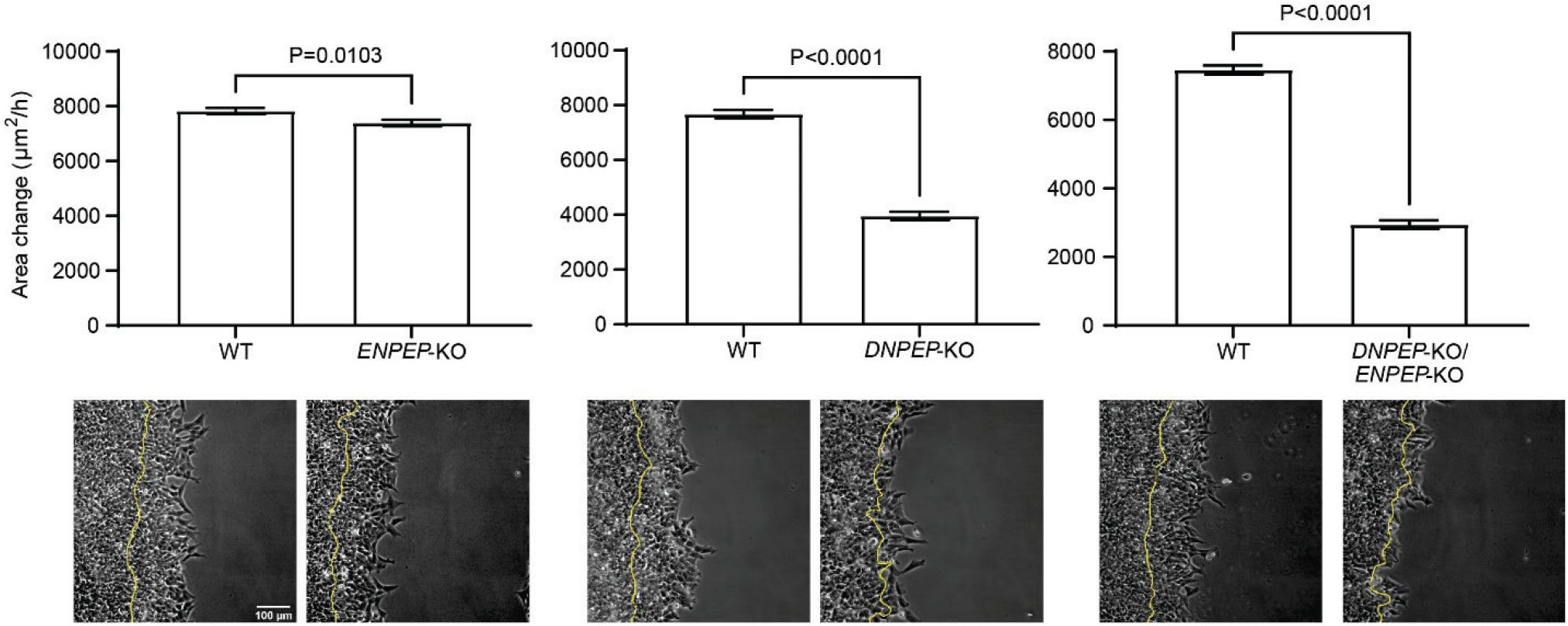
DNPEP knockout affects cell migration. Top, quantification of the cell migration rates in wound healing assays as area covered by the entire wound edge over 12 hours of observation, derived as μm^2^/h; bottom, overlay of phase contrast images of the first (0 h) and last (12 h) frame taken from a representative time lapse videos for each genotype. Yellow line in each image outlines the position of the wound edge at 0 time point. Error bars represent SEM, n= 43 for WT and 44 for ENPEPKO in the left chart, 22 for WT and 19 for DNPEP KO in the middle chart, and 37 for WT and 31 for the double knockout in the right chart.

## Discussion

This study is the first demonstration of previously unknown steps of actin isoform specific N-terminal processing that results in sequential removal of N-terminal Asp residues from beta actin without affecting the closely related gamma actin in the same cells. Actin is one of the striking examples of a protein that possesses a sophisticated enzymatic machinery for multi-step processing of its N-terminus. To date, very few examples of isoform-specific actin processing are known. Our findings add new steps to this processing and further expand the list of isoform-specific actin modifications and potential enzymes involved.

Our results propose two candidate enzymes for mediating N-terminal processing of actin. Aspartyl aminopeptidase (DNPEP) is a cytosolic enzyme that has been largely biochemically characterized, with little insight into its intracellular functions. This enzyme is capable of removing N-terminal Asp or Glu from test substrates structurally similar to synthetic peptides, even though in our in vitro assays it did not target the Asp-containing or Glu-containing actin-based peptides, suggesting that this enzyme’s activity might be generally low, or that it requires additional cellular cofactors in vivo.

ENPEP is largely similar to DNPEP in all these aspects, except that rather than cytosolic, this enzyme is membrane-bound, and it has been previously implicated in regulation of hypertension through the renin-angiotensin system [17]. ENPEP was initially characterized to have a preference for N-terminal Glu-containing substrates, however our in vitro enzymatic assay shows that ENPEP can cleaves both Asp and Glu from the peptides. In principle, both ENPEP and DNPEP should be able to act on any proteins with exposed N-terminal Asp or Glu, however very few of such proteins aside from actin are believed to be naturally produced, since the most common class of aminopeptidases known to remove N-terminal Met from newly synthesized proteins, Met-AP1 and Met-AP2, target largely the proteins in which the initiator Met precedes small and uncharged residues [19]. Actin is a unique protein that possesses a dedicated aminopeptidase to remove its initiator Met to expose a bulky charged residue at the N-terminus [8, 11, 20].

It is surprising that gamma actin, unlike beta actin, never undergoes sequential removal of N-terminal acidic residues, given the fact that both beta and gamma actin N-termini are highly homologous and should both fall within the specificity of Asp and Glu aminopeptidases. Even more puzzling, our results show that this preferential ability of beta, but not gamma actin to undergo sequential removal of N-terminal acidic residues is not a property of the beta actin nucleotide sequence, but is amino acid sequence dependent. In combination, these results suggest that the specificity of this processing toward beta actin is likely not defined by the enzymes responsible for the cleavage itself, but rather by a more complex balance of different intracellular pathways mediating different types of actin processing in vivo. Since gamma actin’s N-terminus is expected to interact stronger with actin N-terminal acetyltransferase NAA80 due to its stronger negative charge [21], it is possible that competition with NAA80 may constitute at least one of the mechanisms regulating this N-terminal processing and its selectivity toward beta actin. Notably, removal of Asp residues from beta actin’s N-terminus is predicted to reduce or abolish its ability to interact with NAA80. It is likely, however, that the interrelationship between these mechanisms is even more complex.

Removal of Asp 2 from the beta actin sequence exposes the beta actin N-terminus for arginylation, previously shown to target N-terminally exposed Asp 3 [12] to affect ∼0.1% of beta actin at a steady state level in HAP1 cells [13]. Notably, deletion of *NAA80* increases this fraction by over 10-fold [5]. This supports the idea that DNPEP and ENPEP may act as additional enzymes that process unacetylated actin to prime it for arginylation, and possibly other modifications or molecular interactions, before the N-terminal acetylation step. Of note, the beta actin species with a single N-terminal Asp appears to be the most prevalent among the processed actin variants, and the only one still present after *ENPEP* knockout. It is possible that a different type of enzyme(s) can remove the Asp-Asp-, Ac-Asp-Asp-, or Met-Asp-Asp di- and tripeptide entities rather than removing single amino acid residues, independently generating this particular actin variant.

Overall, sequential removal of N-terminal beta actin Asp residues targets a very small fraction of intracellular actin, 1-3% or less seen in our studies. It is attractive to suggest that this processing constitutes and additional, previously unknown step in actin regulation – e.g., by exposing actin N-terminus to a different class of molecular interactions. Actin variants with a reduced or missing stretch of negatively charged amino acids at the N-terminus, even if minor, could greatly expand the variety of functions of this already highly multifunctional protein. The questions of the biological mechanisms that lead form this novel type of N-terminal actin processing to actin function in cell migration, cytoskeleton maintenance, filopodia protrusion, and other actin-related processes constitute an exciting direction of future studies.

## Materials and methods

### Cell culture

Human HAP1 cells (wild type, D*NPEP*-KO, *ENPEP*-KO and *DNPEP*/*ENPEP*-KO) were cultured as described by Drazic, A. et al [22]. Immortalized wild type mouse embryonic fibroblasts (MEF) were cultured in DMEM (high glucose with GlutaMAX, Gibco Life Technology) containing 10% fetal bovine serum and 1% Penicillin-Streptomycin (Antibiotic-Antimycotic solution; Life Technology) at 37 °C with 5% CO_2_.

### Vector construction and cell transfection

To generate related gene knockout cells, CRISPR/Csa9 all-in-one expression vectors containing multiple guild RNA expression cassettes and a Cas9 nickase cassette were used for targeted gene deletion (deletion scheme and results were showed in Fig. S4). Detailed procedures were described by Sakuma et al [23]. and the oligos for vector construction and knockout cell genotyping were listed in Table S1. HAP1 cells were cultivated in Iscove’s Modified Dulbecco’s Medium (IMDM) with the addition of 10% FBS and 1% penicillin/streptomycin. Briefly 70-90% confluent cells in 6-well plated were transfected with 1 µg CRISPR/Cas9 plasmids per well by using FuGENE 6 (Promega, Madison, Wisconsin). After 24 h post-transfection, 2 µg/ml puromycin were added to the cell culture and incubated for 48 h followed by splitting cells into 96-well plates to reach 0.5 cell per well.

### Cell migration assays and imaging

An infinite scratch wound was made for cell migration assay. The cells were recovered for 2 h before imaging. Migration rates were measured as the area covered by the edge of the wound in the field of view per unit time using Fiji. Images were acquired by a Nikon Ti microscope with 20X Phase objective and Andor iXon Ultra 888 EMCCD camera.

### Immunofluorescence staining

To quantitate the amount of F-actin, spreading area and filopodia, HAP1 cell culture with 70-80% confluence were trypsinized and diluted 4 fold followed by seeded on coverslips in 6-well plates. The cells were cultivated for 24 h and then fixed in 4% paraformaldehyde aqueous solution (PFA) at room temperature for 30 min followed by washing with PBS three times. Cells were then stained with Phalloidin conjugated to AlexaFluor 594 (Molecular Probes, Eugene, OR) and DAPI (Thermo Fisher Scientific, Waltham, Massachusetts). Cell images for F-actin quantification were acquired by using Nikon Ti microscope with 100X objective and Andor iXon Ultra 888 EMCCD camera. Cell filopodia were imaged by Nikon Ti microscope with 100X objective lens, equipped with a spinning disk CSU-22 (Yokogawa) and Andor iXon Life IXON-L-897 EMCCD camera. All images were analyzed by fiji (NIH, Bethesda, MD).

### Protein and synthetic peptide preparation for mass spectrometry

HAP1 and MEF cells with 70-80% confluence were harvested by scraping and centrifugation. The pellets were washed by PBS, and spun down to remove the supernatant. The cell pellets were lysed directly in 4 × SDS sample buffer at the W:V ratio of 1: 10 (1 mg cell : 10 μL buffer), followed by boiling in water for 10 min. 10 μL of each sample was loaded for SDS-PAGE electrophoresis at 150 V. The actin bands were excised from the gel, reduced with tris(2-carboxyethyl)phosphine, alkylated with iodoacetamide, and digested with trypsin. The trypsin digested actin were analyzed using a standard 90-min LC gradient on the Thermo Q Exactive Plus mass spectrometer. The mass spectrometry data were searched with partial tryptic specificity against the mammalian actin isoform database and a contaminant database using MaxQuant 1.6.17.0 [24].

The synthetic peptides (Ac-DDDIAALC, Ac-EEEIAALC, DDDIAALC, EEEIAALC) were synthesized by GenScript (Piscataway, NJ). The DNPEP and ENPEP proteins were purchased from OriGene (Cat#: TP300104) and Novus Biologicals (Cat#: 2499-ZN-010) repectively. The enzymatic assay was modified from ENPEP assay procedures: 200 µL reaction system (25 mM Tris, 50 mM CaCl2, 0.2 M NaCl, pH 8.0) containing 100 µM peptide and 400 ng recombinant proteins were incubated at 37 ºC for 30 min, and then the reactions were terminated by heating at 95°C for 15 min followed by incubation of the tubes on ice for 20 min. The tubes were then centrifuged at 17,000 x g for 15 min at RT and the supernatants were loaded onto C18 spin columns prewashed with 100% acetonitrile and water by centrifugation at 110 x g for 1 min. The columns were washed twice with 150 µL peptide wash buffer (0.1% trifluoroacetic acid (TFA) in water), followed by elution of the column-bound peptides with 150 µL of peptide elution buffer, repeated two times to ensure the complete peptide recovery. The eluates were concentrated to approximately 20 µL using a SpeedVac vacuum concentrator. 5 µL of 20 mM TCEP was added to 10 µL of sample and incubated for 30 min at 37 ºC. The samples were analyzed on the Thermo Q Exactive HF mass spectrometer. Data were analyzed using the Thermo Xcalibur software.

### Identification and Quantification of N-terminal processing of actin isoforms

Peptide sequences were identified using MaxQuant 1.6.17.0 [24]. MS/MS spectra were searched against a custom database containing only the mammalian actin sequences, using partial tryptic specificity, static carboxamidomethylation of Cys, and variable Met oxidation, protein N-terminal acetylation, and Asn deamidation.

For quantification of the actin N-terminal peptides, all the peptides corresponding to the beta and gamma actin N-termini were separated out and divided into groups according to their N-terminal residue (Met1, acetylated Asp/Glu2, Asp/Glu3, Asp/Glu4, or Ile/Val5), regardless of any additional variable modifications. Of note, no N-terminal acetylation on Met or the residues past the Asp/Glu2 were detected. Intensities of each peptide species were added together and divided by the total intensity added for the entire repertoire of the N-termini for each actin isoform to obtain fractions of the total shown in the figures.

## Acknowledgements

We thank Dr. Pavan Vedula for helpful discussions and Thomas Beer from the Wistar Institute Proteomics Facility for the analysis of in vitro N-terminally processed peptides. This work was supported by the NIH grant R35GM122505 to AK and R50 CA221838 to HYT.

## References

1. Bunnell, T.M. and J.M. Ervasti, Delayed embryonic development and impaired cell growth and survival in Actg1 null mice. Cytoskeleton (Hoboken), 2010. 67(9): p. 564–72.

2. Vedula, P. and A. Kashina, The makings of the ‘actin code’: regulation of actin’s biological function at the amino acid and nucleotide level. J Cell Sci, 2018. 131(9).

3. Bunnell, T.M. and J.M. Ervasti, Structural and functional properties of the actin gene family. Crit Rev Eukaryot Gene Expr, 2011. 21(3): p. 255–66.

4. Bunnell, T.M., et al., beta-Actin specifically controls cell growth, migration, and the G-actin pool. Mol Biol Cell, 2011. 22(21): p. 4047–58.

5. Vedula, P., et al., Diverse functions of homologous actin isoforms are defined by their nucleotide, rather than their amino acid sequence. Elife, 2017. 6.

6. Vedula, P., et al., Different translation dynamics of beta- and gamma-actin regulates cell migration. Elife, 2021. 10.

7. Rubenstein, P.A. and D.J. Martin, NH2-terminal processing of actin in mouse L-cells in vivo. J Biol Chem, 1983. 258(6): p. 3961–6.

8. Solomon, L.R. and P.A. Rubenstein, Correct NH2-terminal processing of cardiac muscle alpha-isoactin (class II) in a nonmuscle mouse cell. J Biol Chem, 1985. 260(12): p. 7659–64.

9. Drazic, A., et al., NAA80 is actin’s N-terminal acetyltransferase and regulates cytoskeleton assembly and cell motility. Proc Natl Acad Sci U S A, 2018. 115(17): p. 4399–4404.

10. Terman, J.R. and A. Kashina, Post-translational modification and regulation of actin. Curr Opin Cell Biol, 2013. 25(1): p. 30–8.

11. MacTaggart, B. and A. Kashina, Posttranslational modifications of the cytoskeleton. Cytoskeleton (Hoboken), 2021. 78(4): p. 142–173.

12. Karakozova, M., et al., Arginylation of beta-actin regulates actin cytoskeleton and cell motility. Science, 2006. 313(5784): p. 192–6.

13. Chen, L. and A. Kashina, Quantification of intracellular N-terminal beta-actin arginylation. Sci Rep, 2019. 9(1): p. 16669.

14. Zhang, F., et al., Differential arginylation of actin isoforms is regulated by coding sequence-dependent degradation. Science, 2010. 329(5998): p. 1534–7.

15. Carette, J.E., et al., Ebola virus entry requires the cholesterol transporter Niemann-Pick C1. Nature, 2011. 477(7364): p. 340–3.

16. Wilk, S., E. Wilk, and R.P. Magnusson, Purification, characterization, and cloning of a cytosolic aspartyl aminopeptidase. J Biol Chem, 1998. 273(26): p. 15961–70.

17. Yang, Y., et al., Structural insights into central hypertension regulation by human aminopeptidase A. J Biol Chem, 2013. 288(35): p. 25638–25645.

18. Perrin, B.J. and J.M. Ervasti, The actin gene family: function follows isoform. Cytoskeleton (Hoboken), 2010. 67(10): p. 630–4.

19. Li, X. and Y.H. Chang, Amino-terminal protein processing in Saccharomyces cerevisiae is an essential function that requires two distinct methionine aminopeptidases. Proc Natl Acad Sci U S A, 1995. 92(26): p. 12357–61.

20. Chen, L. and A. Kashina, Post-translational Modifications of the Protein Termini. Front Cell Dev Biol, 2021. 9: p. 719590.

21. Rebowski, G., et al., Mechanism of actin N-terminal acetylation. Sci Adv, 2020. 6(15): p. eaay8793.

22. Drazic, A., et al., NAA80 is actin’s N-terminal acetyltransferase and regulates cytoskeleton assembly and cell motility. Proceedings of the National Academy of Sciences, 2018. 115(17): p. 4399–4404.

23. Sakuma, T., et al., Multiplex genome engineering in human cells using all-in-one CRISPR/Cas9 vector system. Scientific reports, 2014. 4.

24. Cox, J. and M. Mann, MaxQuant enables high peptide identification rates, individualized p.p.b.-range mass accuracies and proteome-wide protein quantification. Nat Biotechnol, 2008. 26(12): p. 1367–72.

