## Supplemental Figures for "Differential N-terminal processing of beta and gamma actin in vivo"

Supplemental Online Information

Table S1: Primers used for CRISPR/Cas9 gene editing and genotyping

| Primer name | Sequence (5’-3’) |
|---|---|
| DNPEP-guide RNA-1-F | caccgTGTCCGTGTGGGCCCCGATG |
| DNPEP-guide RNA-1-R | aaacCATCGGGGCCCACACGGACAc |
| DNPEP-guide RNA-2-F | caccgTAAGGAACTCGGACGGTGTT |
| DNPEP-guide RNA-2-R | aaacAACACCGTCCGAGTTCCTTAc |
| DNPEP-guide RNA-3-F | caccgTCTCTTACCACAGCATTGAG |
| DNPEP-guide RNA-3-R | aaacCTCAATGCTGTGGTAAGAGAc |
| DNPEP-guide RNA-4-F | caccgTCTCCTGATCATGGGGAGGG |
| DNPEP-guide RNA-4-R | aaacCCCTCCCCATGATCAGGAGAc |
| DNPEP-genotyping primer-F | TGAACGGTAAGGCCCGCAAAGA |
| DNPEP-genotyping primer-R | CGTGTTCAAGCAGGACACAGCA |
| ENPEP-guide RNA-1-F | caccGAAACAAGGGGGGTGGAATG |
| ENPEP-guide RNA-1-R | aaacCATTCCACCCCCCTTGTTTC |
| ENPEP-guide RNA-2-F | caccGGCTGGTGGGAACGTGAAAA |
| ENPEP-guide RNA-2-R | aaacTTTTCACGTTCCCACCAGCC |
| ENPEP-guide RNA-3-F | caccGAGGGAGCCGTTCAGCCAGC |
| ENPEP-guide RNA-3-R | aaacGCTGGCTGAACGGCTCCCTC |
| ENPEP-guide RNA-4-F | caccgATTTTATAGAACCACCTACA |
| ENPEP-guide RNA-4-R | aaacTGTAGGTGGTTCTATAAAATc |
| ENPEP-genotyping-F | TTCCTGAGCTTGTCATTCAGAAACA |
| ENPEP-genotyping-R | CATGAGCAACAAGCCTCCAA |

Small caps: for BbSI enzymatic digested plasmid recognition; red small caps: G is added to the 5’ position for efficient U6 promoter transcription.

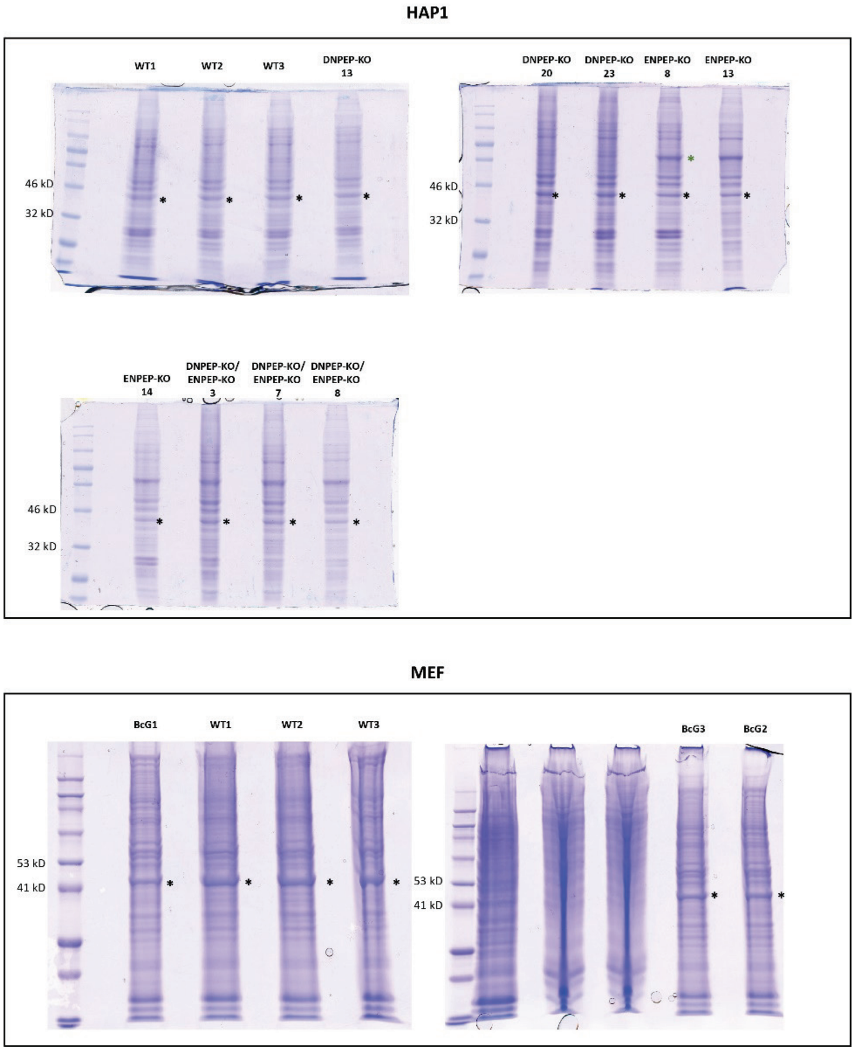


Figure S1. Images of SDS polyacrylamide gels used for the excision and analysis of the actin gel bands. Stars indicate the band cut from the gel and used for mass spectrometry analysis.

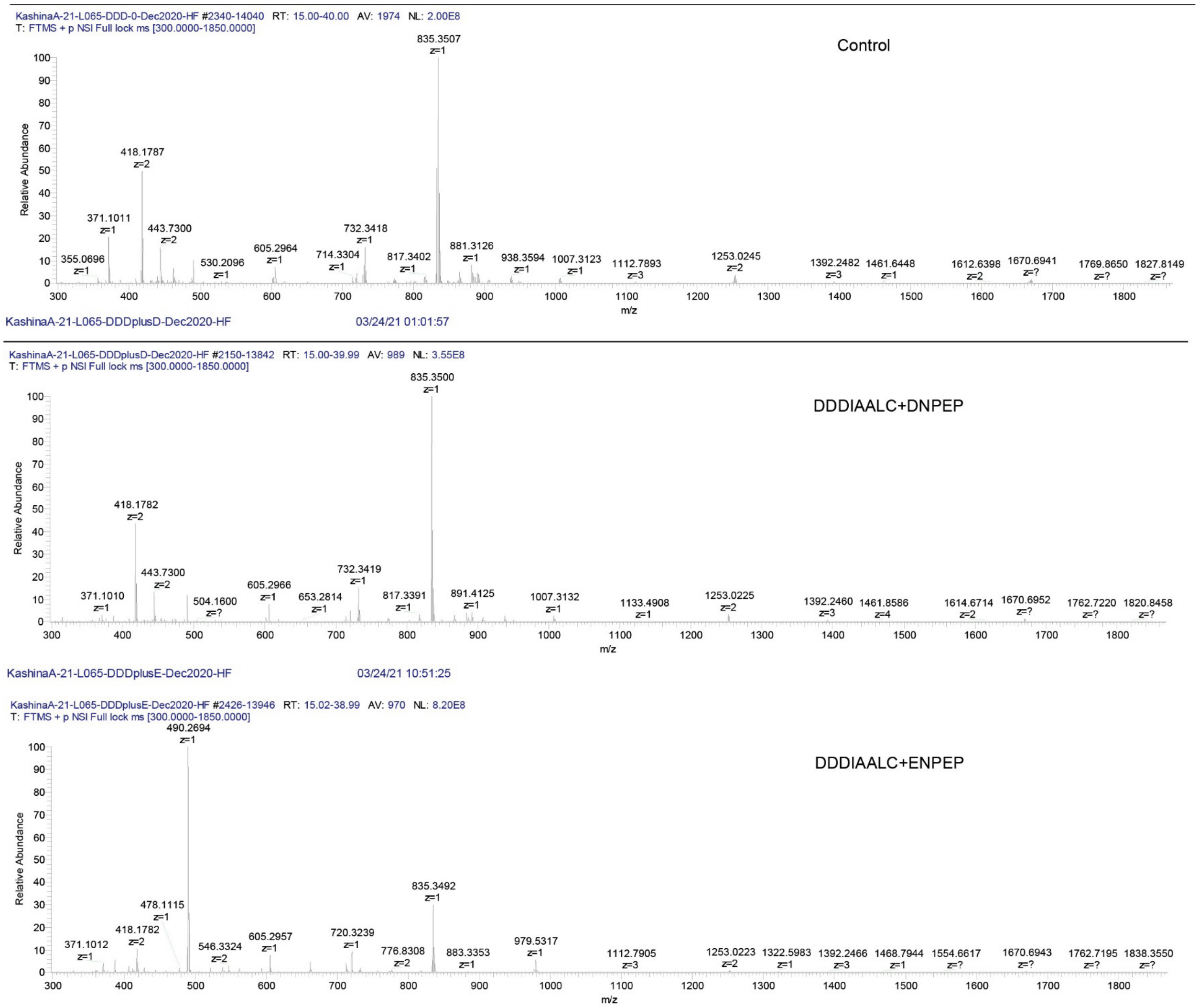


Figure S2A. Mass spectrometry chromatograms of unacetylated peptides corresponding to the beta actin N-terminus, before and after addition of DNPEP or ENPEP

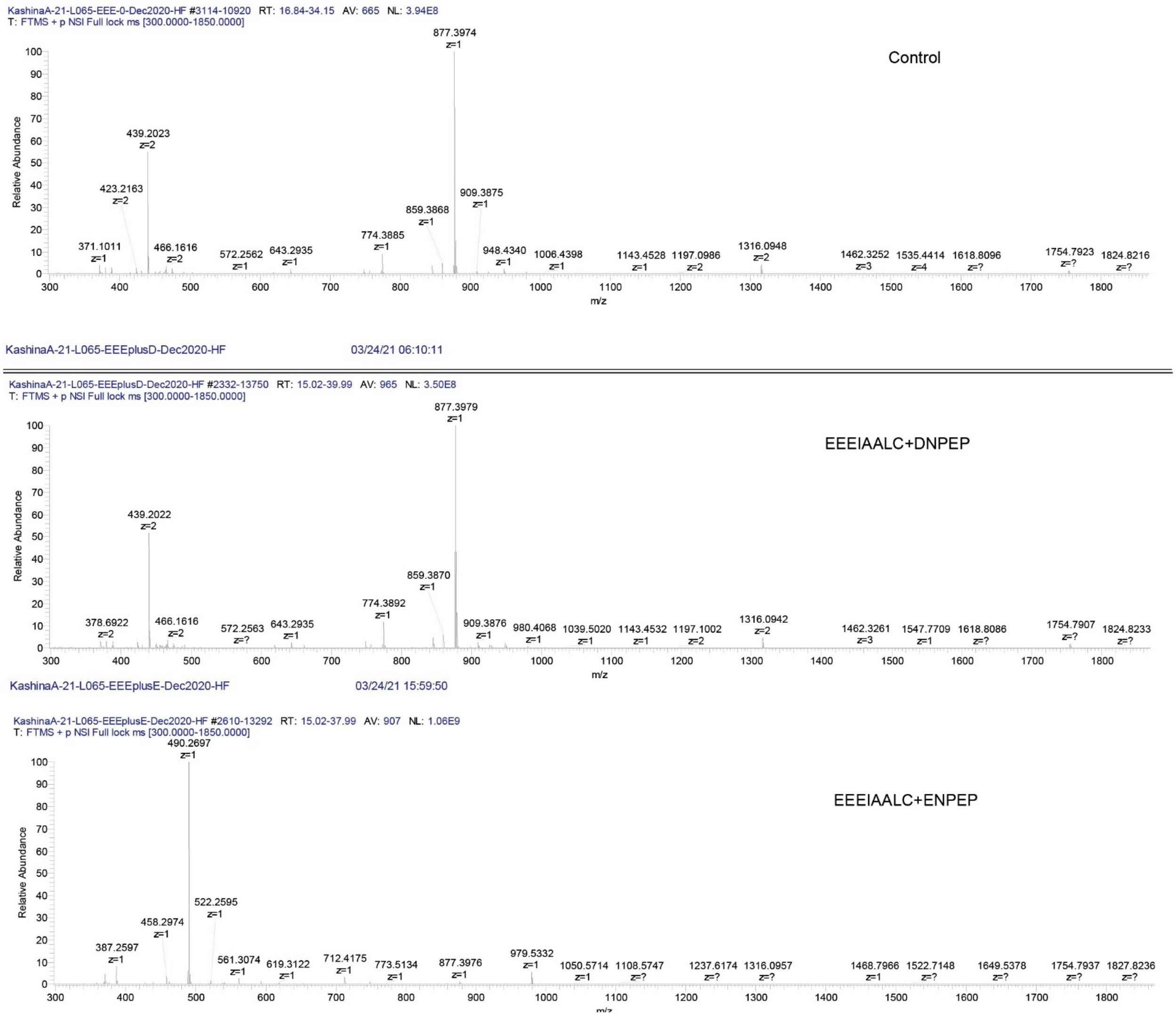


Figure S2B. Mass spectrometry chromatograms of unacetylated peptides corresponding to the gamma actin N-terminus, before and after addition of DNPEP or ENPEP

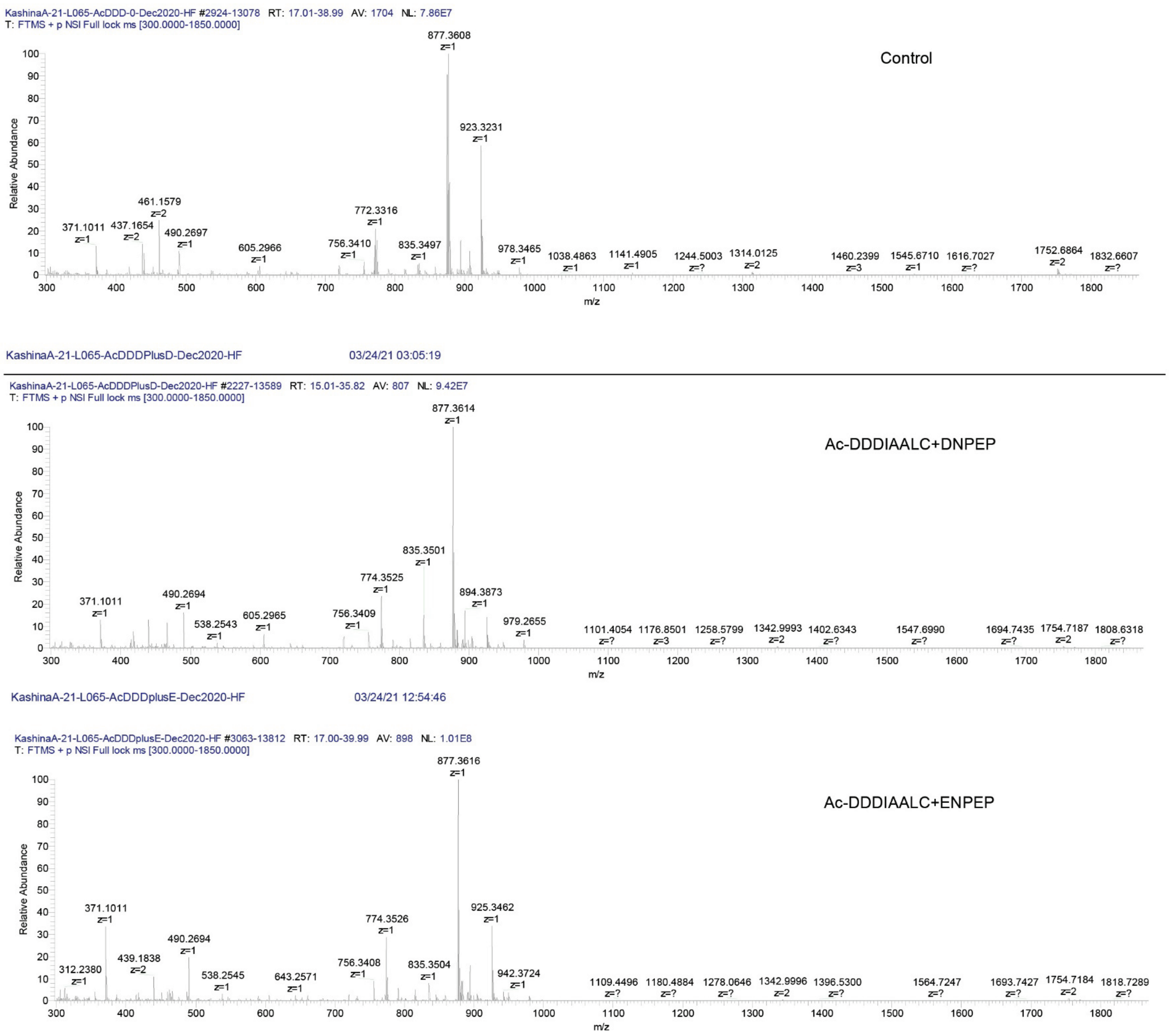


Figure S3A. Mass spectrometry chromatograms of acetylated peptides corresponding to the beta actin N-terminus, before and after addition of DNPEP or ENPEP

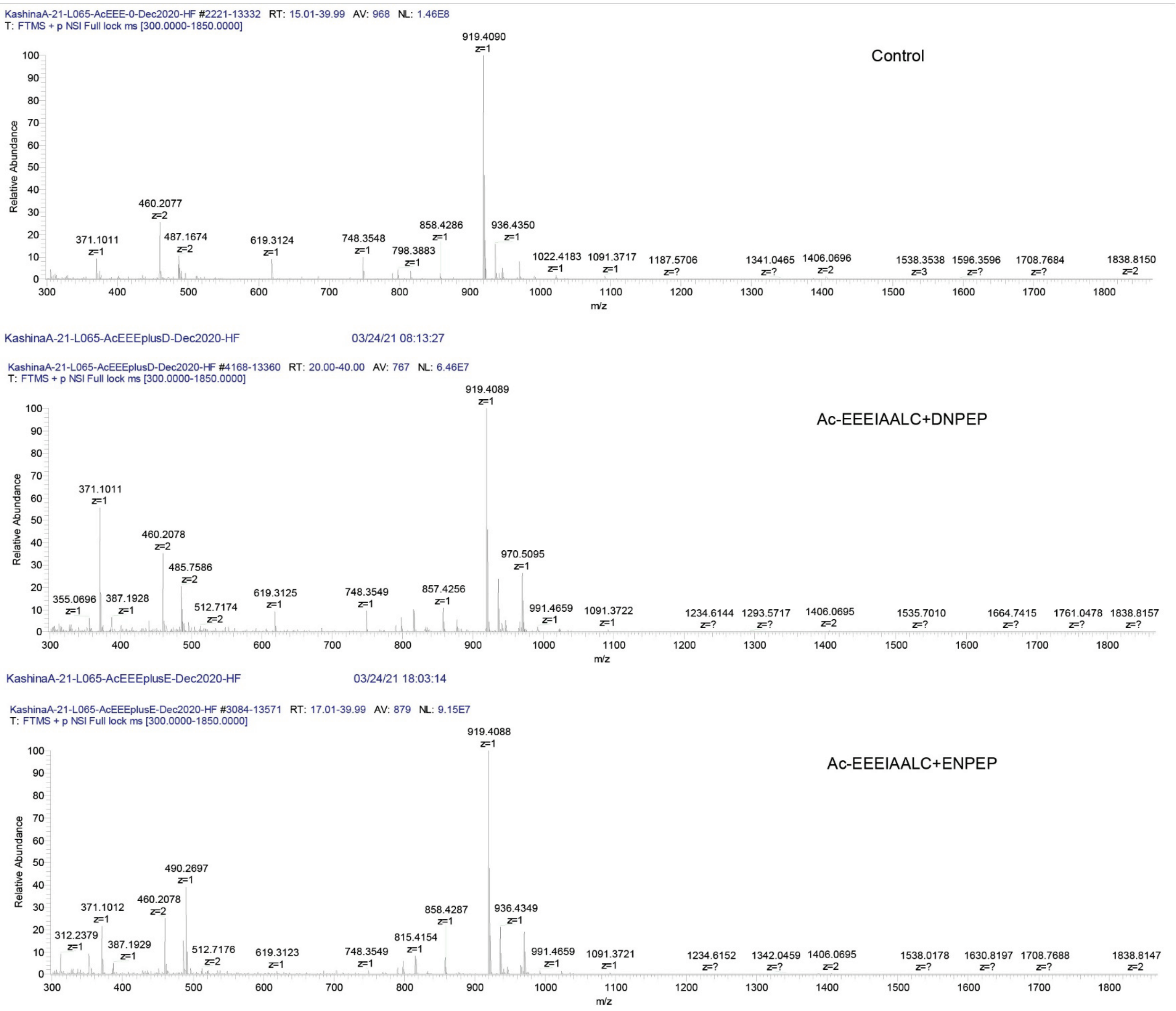


Figure S3B. Mass spectrometry chromatograms of acetylated peptides corresponding to the gamma actin N-terminus, before and after addition of DNPEP or ENPEP

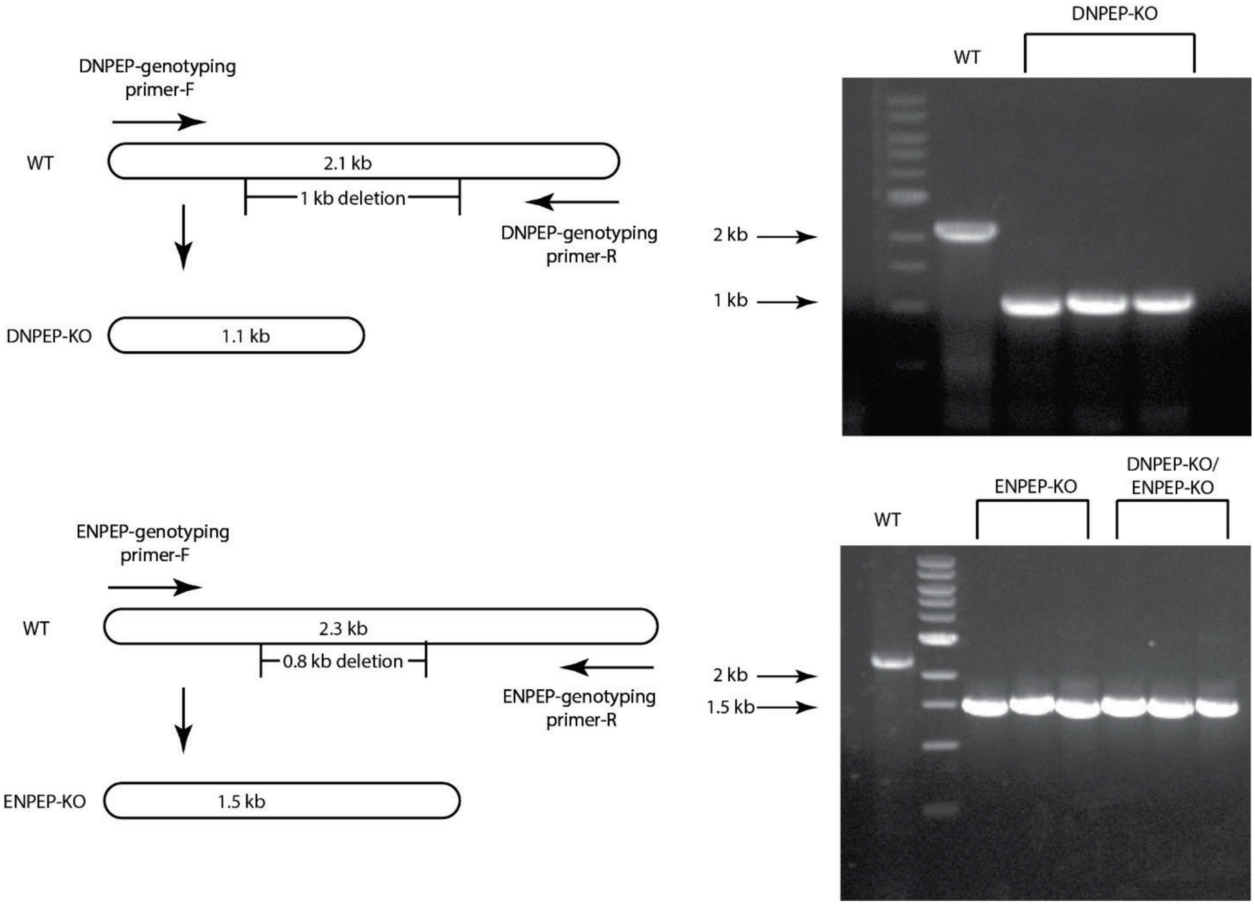


Figure S4. Schematic illustration of gene knockout strategy and genotyping (left), and representative genotyping gels (right)
